# Interleukin-6 Receptor Signalling and Abdominal Aortic Aneurysm Growth Rates

**DOI:** 10.1101/428516

**Authors:** Ellie Paige, Marc Clément, Fabien Lareyre, Michael Sweeting, Juliette Raffort, Céline Grenier, Alison Finigan, James Harrison, James E. Peters, Benjamin B. Sun, Adam S. Butterworth, Seamus C. Harrison, Matthew J. Bown, Jes S. Lindholt, Stephen A. Badger, Iftikhar J. Kullo, Janet Powell, Paul E. Norman, D Julian A. Scott, Marc A. Bailey, Stefan Rose-John, John Danesh, Daniel F. Freitag, Dirk S. Paul, Ziad Mallat

**Affiliations:** National Centre for Epidemiology and Population Health, Research School of Population Health, The Australian National University, Canberra, Australia; Division of Cardiovascular Medicine, University of Cambridge, Cambridge, UK; Université Côte d’Azur, Institut National de la Sante et de la Recherche Medicale, Centre Mediterranéen de Recherche Moleculaire, Nice, France; University Hospital of Nice, France; BHF Cardiovascular Epidemiology Unit, Department of Public Health and Primary Care, University of Cambridge, Cambridge, UK; Department of Health Sciences, University of Leicester, Leicester, UK; British Heart Foundation Centre of Excellence, University of Cambridge School of Clinical Medicine, Cambridge, UK; NIHR Blood and Transplant Research Unit in Donor Health and Genomics, Cambridge, UK; Department of Cardiovascular Sciences and NIHR Leicester Biomedical Research Centre, University of Leicester, Leicester, UK; Department of Cardiovascular and Thoracic surgery, Elitary Reserach Centre of Individualised Medicine in Arterial Disease, Odense University Hospital, Odense, Denmark; Regional Vascular Surgery Unit, Belfast Health & Social Care Trust, Belfast, UK; Department of Cardiovascular Medicine and the Gonda Vascular Center, Mayo Clinic, Rochester, Minnesota, USA; Faculty of Medicine, Department of Surgery and Cancer, Imperial College London, London, UK; Medical School, University of Western Australia, Perth, Australia; Leeds Vascular Institute, Leeds General Infirmary, Leeds, UK; Leeds Institute of Cardiovascular & Metabolic Medicine, School of Medicine, University of Leeds, Leeds, UK; Department of Biochemistry, Christian-Albrechts-University, Kiel, Germany; Department of Human Genetics, Wellcome Sanger Institute, Hinxton, UK; Institut National de la Santé et de la Recherche Médicale, Paris Cardiovascular Research Center, France

**Author notes:** Joint first authors (Ellie Paige, Marc Clément). Joint senior authors (Daniel F. Freitag, Dirk S. Paul, Ziad Mallat). Correspondence: Professor Ziad Mallat, Department of Medicine, University of Cambridge, Cambridge, CB2 0QQ, UK.

**Keywords:** Animal Models of Human Disease, Inflammation, Epidemiology, Genetic, Association Studies, Aneurysm

## Abstract

**Background:** The Asp358Ala variant (rs2228145; A>C) in the interleukin-6 receptor (*IL6R*) gene has been implicated in the development of abdominal aortic aneurysms (AAAs), but its effect on AAA growth over time is not known. We aimed to investigate the clinical association between the *IL6R*-Asp358Ala variant and AAA growth, and to assess the effect of blocking the IL-6 signalling pathway in mouse models of aneurysm rupture.

**Method:** Using data from 2,863 participants with AAA from nine prospective cohorts, age- and sex-adjusted mixed-effects linear regression models were used to estimate the association between the *IL6R*-Asp358Ala variant and annual change in AAA diameter (mm/year). In a series of complementary randomised trials in mice, the effect of blocking the IL-6 signalling pathways was assessed on plasma biomarkers, systolic blood pressure, aneurysm diameter and time to aortic rupture and death.

**Results:** After adjusting for age and sex, baseline aneurysm size was 0.55mm (95% confidence interval [CI]: 0.13, 0.98mm) smaller per copy of the minor allele [C] of the Asp358Ala variant. There was no evidence of a reduction in AAA growth rate (change in growth=-0.06mm per year [−0.18, 0.06] per copy of the minor allele). In two mouse models of AAA, selective blockage of the IL-6 trans-signalling pathway, but not combined blockage of both, the classical and trans-signalling pathways, was associated with improved survival (p<0.05).

**Conclusions:** Our proof-of-principle data are compatible with the concept that IL-6 trans-signalling is relevant to AAA growth, encouraging larger-scale evaluation of this hypothesis.

## Introduction

Abdominal aortic aneurysms (AAAs) are defined as an enlargement of the aorta to ≥30mm diameter. They usually grow asymptomatically until rupture occurs, after which the survival of affected individuals is less than 20%.^1^ AAAs typically occur in mid-to-later life and more commonly in men (prevalence of 1-2%^2, 3^ compared to <1% in women^4^). Current standard of care is surgical intervention, either open surgery or endovascular repair. However, due to surgical risks vs. benefits, such interventions are generally recommended only for people with larger AAAs (diameter ≥55mm or >40mm and enlarging >10mm/year).^5, 6^ The growth rates of AAAs vary considerably between individuals^7^ and there is currently a lack of therapeutic options to slow or halt progression of AAAs.^8^ Inflammatory processes in the vessel wall may contribute to the progression of AAAs.^9-11^ For example, levels of circulating inflammatory markers including interleukin (IL)-6 are higher in prevalent AAA cases than controls^12^ and correlate with the size of AAA in cross-sectional studies.^13^

IL-6 is a central coordinator of inflammatory responses by controlling the systemic inflammatory response in the liver and the activation and differentiation of leukocyte subsets, including macrophages and T cells. IL-6 signalling occurs in two different modes, termed “classical” (cis-) and trans-signalling. In classical signalling, binding of IL-6 to the membrane-bound IL-6 receptor (mIL-6R) induces homodimerization with its co-receptor gp130, resulting in the phosphorylation of the transcription factors STAT3 and STAT1.^14, 15^ Hence, classical signalling is dependent on the membrane-bound form of the IL-6 receptor and occurs only in leukocyte subsets and hepatocytes that express this molecule. By contrast, trans-signalling occurs through a circulating soluble form of IL-6R (sIL-6R), which, if bound to IL-6, is able to stimulate cells expressing gp130, even in the absence of mIL-6R.^14, 16^ Because gp130 is almost ubiquitously expressed, IL-6 trans-signalling can occur in virtually any cell, although is probably active only during conditions of immunological stress.^14, 16^

A non-synonymous variant (Asp358Ala; rs2228145 A>C) in the *IL6R* gene, encoding the IL-6 receptor, plays a critical role in IL-6 signalling. The minor allele [C] of this variant is associated with a reduced risk of several chronic conditions including coronary heart disease,^17^ atrial fibrillation,^18^ rheumatoid arthritis,^19^ type 1 diabetes,^20^ but an increased risk of asthma.^21^ The variant results in more efficient proteolytic cleavage of mIL-6R, thereby reducing levels of mIL-6R and dampening classical signalling.^22, 23^ Conversely, the variant increases levels of sIL-6R, although the exact effects of the variant on the trans-signalling pathway are unknown.^20^ A Mendelian randomisation study has previously implicated the rs2228145 in the causal pathway of AAA, with the minor allele [C] showing a protective effect for the risk of AAA and a combined endpoint of rupture or surgical intervention.^12^ Licenced drugs are available that target the IL-6/IL-6 receptor pathway. However, evidence is needed that this pathway is associated with aneurysm progression or rupture in order to encourage repurposing drugs for use in patients with known AAA.

The aims of this study were to: (1) to assess and quantify the effect of the functional *IL6R* variant on the progression of AAAs in population cohorts with prospective follow-up and standardized repeated measurements of AAA diameter; and (2) estimate the effect of blocking the IL-6 signalling pathway (i.e., either both classical and trans-signalling pathways or specifically the trans-signalling pathway) on time to aneurysm rupture in mouse models.

## Material and methods

### Human genetic analyses

#### Study participants

We analysed previously collected data from six prospective cohorts contributing to the RESCAN collaboration and three others (Belfast, Mayo Clinic [New York] and St George’s [London]), that had repeated measurements of aneurysm diameter over time and genotyped data either for the *IL6R*-Asp358Ala variant (rs2228145) or for one of two proxy variants (rs4129267 or rs7529229; both r^2^=1.0 with rs2228145, 1000 Genomes project). Genotype data for these cohorts was provided by the Aneurysm Consortium. Details on the RESCAN collaboration have been previously published and a list of RESCAN and Aneurysm Consortium collaborators is provided in Appendices 1 and 2. AAA was defined as having an aorta size ≥30mm.^1^ Aneurysm diameter were measured by ultrasound, CT scan or MRI. Only participants that met the definition for AAA and had at least two diameter measurements on different dates were included in the current study. Each cohort was approved by a research ethics committee and all participants gave informed consent.

#### Statistical analyses

Age- and sex-adjusted mixed-effects linear regression models were used to first model the change in AAA growth over time and then to estimate the association between rs2228145 (or the proxy variants rs4129267 or rs7529229) and baseline AAA size and annual change in AAA diameter (mm/year). The main analysis was pre-specified to test the hypothesis that rs2228145 is associated with a decreased rate of AAA growth. Effect estimates and standard errors were pooled using inverse-variance weighted fixed-effects meta-analysis. 95% confidence intervals (CIs) were calculated for all effect sizes and an overall p-value for the pooled effect was calculated. Heterogeneity between studies was quantified using the I^2^ statistic.^24^ To test whether adjustment for additional potential confounders changed the effect size, we re-ran the main analysis in a sub-sample of six studies with recorded data on current smoking, diabetes status, body mass index and method used to measure aneurysm size (i.e., ultrasound, CT scan, MRI or not specified), adjusting for these additional covariates.

#### Sensitivity analyses

Four sensitivity analyses were conducted. First, since participants with initially large aneurysms (i.e., those above the surgery threshold of 55mm) might be categorically different to participants with smaller aneurysms,^25^ we re-ran the main model examining annual change in AAA diameter, and restricted the analysis to participants with a small baseline aneurysm diameter of 30-44mm. Aneurysms that grew larger than 45mm were censored in the analysis after the first measurement ≥45mm. Second, we also re-ran the main model restricting the analysis to participants with a medium baseline aneurysm diameter of 45-54 mm. Aneurysms that grew larger than 55mm were censored in the analysis after the first measurement ≥55mm. Third, a Cox proportional hazards model was used to examine the association between the rs2228145 variant (or proxy variant) and the rate of reaching the surgery threshold of ≥55mm, with baseline hazards estimated within 5mm baseline aneurysm size groups. Fourth, to increase statistical power, we re-ran the main analysis including participants with a single diameter AAA measure. The sensitivity analyses were exploratory and not pre-specified.

Power calculations for the main analysis are provided in Appendix 3. All analyses were done using Stata 13.1.

#### Phenome-scan

To explore the mechanisms through which inhibition of IL6-R signalling could influence AAA, we performed a genetic association analysis of the rs2228145 variant with a large panel of potentially relevant intermediate traits and cytokine profiles in healthy participants across several human genetic studies. We collected genetic association data for 36 white cell, red cell, and platelet properties in up 173,039 European-ancestry individuals;^26^ systolic/ diastolic blood pressure and pulse pressure in up to 161,871 individuals;^27^ and a wide range of plasma proteins in up to 4,981 individuals.^28^

### Experimental mouse models

#### Mouse models

Several mouse models have been established that reproduce certain characteristics of aortic aneurysm development in humans, including inflammation, destruction of the extracellular matrix and aortic dilatation, but all have limitations with regards to the pathological interpretation of human AAA.^29^ We tested the effect of blocking the IL-6 pathway on aortic rupture using two distinct experimental mouse models to study the role of IL-6 signalling in dissecting and non-dissecting AAAs. To investigate dissecting AAAs, mice were treated with a systematic infusion of angiotensin II with pharmacological inhibition of transforming growth factor-β (TGFβ) activity.^29^ To examine non-dissecting AAAs, elastase was applied on top of the aorta and TGFβ activity was inhibited pharmacologically.^30^ The angiotensin II and elastase models are widely used models for AAA, which are characterised by frequent ruptures and high reproducibility.^29^ Blockage of the activity of TGFβ, a critical factor that maintains aortic wall integrity, exacerbates aneurysmal aortic dilatation, induces intraluminal thrombus formation and promotes aortic rupture, thereby reproducing the main features of human AAA.^30^ Details on the development and characterisation of these mouse models have been previously published.^29, 30^

#### Procedures

For all experiments, we used male, C57Bl/6J, 8-week-old mice (Charles River, UK). Mice were infused with angiotensin II (Sigma) at a rate of 1μg/min/kg using osmotic minipumps (model 2004; ALZET). Alternatively, mice underwent surgery for the application of elastase (E1250; Sigma) at 10μl of the filtered solution for 3 minutes on top of the infrarenal aorta. Mice were injected intraperitoneally with anti-TGFβ (clone 1D11; BioXcell) using 250 μg/mouse at 3 times/week, starting on the day of the surgery (minipump implantation or application of elastase). All experiments were ended when the mortality reached 80%. Necropsies were performed to confirm the aortic rupture (thoracic or retroperitoneal hematoma) and aortic tissue samples were harvested, fixed in PFA 4 % overnight at 4°C, and stored in PBS at 4°C until further investigations. The analysis of the tissue sample was performed on all the animals, regardless of the presence of a rupture.

In both mouse models, mice were injected intravenously with a first bolus of 2mg of anti-IL6-R (clone MR16-1^31^; Chugai Pharmaceutical) or control isotype (clone HPRN; Bioxcell) one week before the experiment in order to induce immune tolerance to the antibody.^32^ After surgery, mice were injected intraperitoneally with 500μg of anti-IL-6R twice a week to completely block the IL-6 pathway (i.e. inhibiting both classical and trans-signalling pathways) or with 500μg of a control isotype twice a week. We measured IL-6 concentration in the plasma of mice to confirm that the IL-6 pathway was blocked in the mice treated with anti-IL-6R. In a separate experiment, the IL-6 trans-signalling pathway was selectively blocked using the fusion protein sgp130Fc^33^ (supplied by Prof Stefan Rose-John, Institute of Biochemistry, Christian-Albrechts-University of Kiel, Germany). The mice received 10μg of sgp130Fc or control isotype (human IgG1 Fc; Bioxcell) three times a week starting from the first day of the experiment.

#### Phenotyping

Outcome measures included plasma levels of different biomarkers, systolic blood pressure, recruitment of T cells (CD3 staining) and neutrophils (MPO or Ly6G staining) to the aortic wall, collagen content of the aortic wall (Sirius red staining), aneurysm diameter, and time to aortic rupture and death. The experimental procedures are described in detail in Appendix 4.

Animal experiments were approved by the UK Home Office and performed under PPL PA4BDF775. The care and use of all mice in this study was carried out in accordance with the UK Home Office regulations under the Animals (Scientific Procedures) Act 1986.

## Results

### Association of IL6R-Asp358Ala with AAA growth rate

We studied a total of 2,863 participants across nine prospective cohorts, 91% of whom were men (mean age of 72 years; Table 1). AAA size was on average 43mm at baseline and participants had an average of four scans across an average of three years (Table 1). A summary of baseline characteristics by *IL6R*-rs2228145 (or proxy) genotype is given in Supplementary Table 1. On average, aneurysms grew by 1.88mm per year (95% CI: 1.79, 1.96; Supplementary Figure 1). Baseline aneurysm size was 0.55mm (0.13, 0.98) smaller per copy of the minor allele [C] (Supplementary Figure 2). Among those with at least two measurements of aneurysm size (n=2,154), there was no statistically significant decrease in AAA growth rate (growth per minor allele= −0.06mm/year [−0.18, 0.06]) (Figure 1). All association analyses were adjusted for age and sex. Results were similar after adjustment for current smoking status, diabetes status, body mass index and aneurysm measurement method (Supplementary Figure 3).

**Figure 1:**
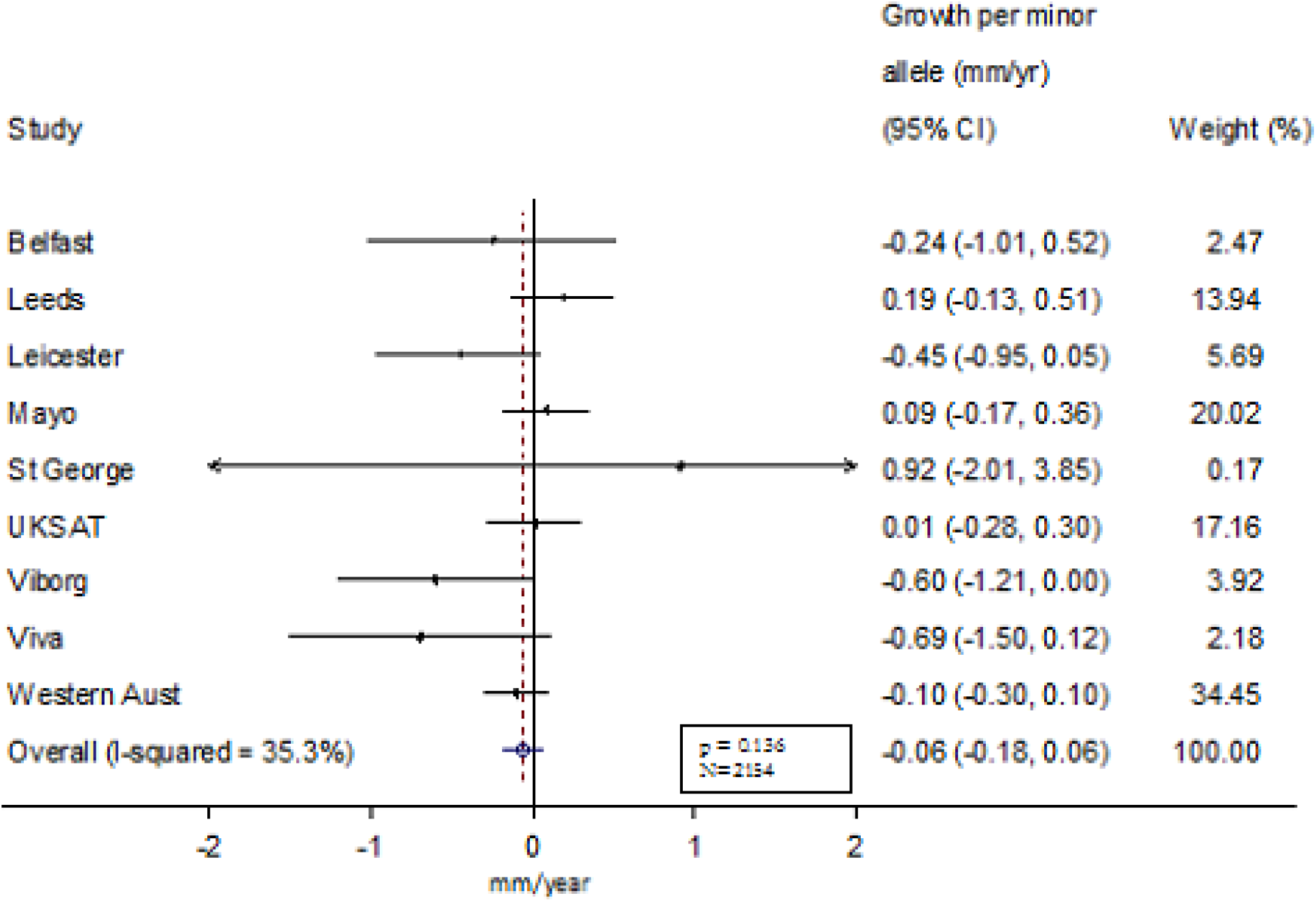
Sex- and age-adjListed change in abdominal aortic an eurysm growth rate (mm/year) per copy of the IL-6 minor allele.

**Table 1.**
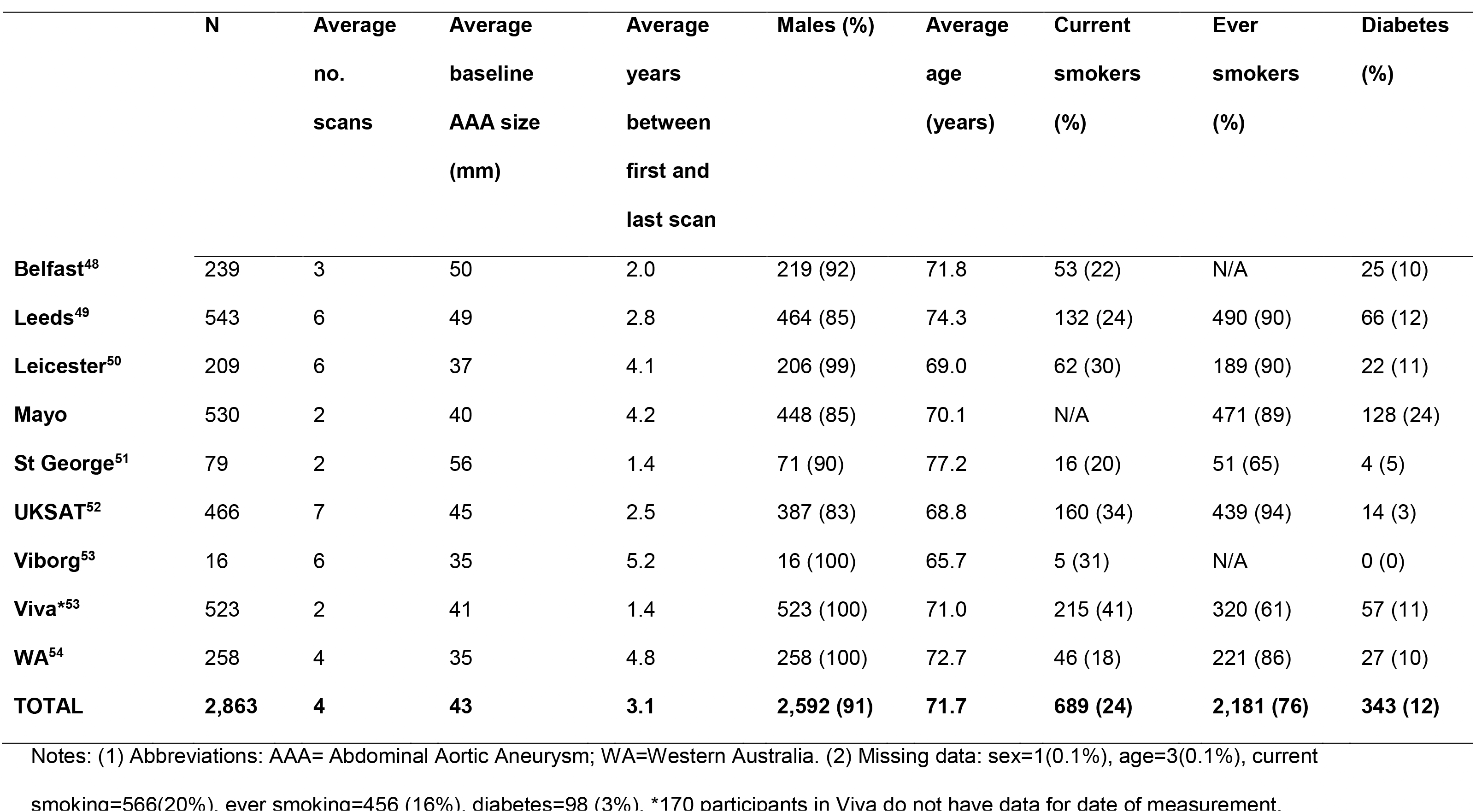
Baseline characteristics of the studies included in the human genetic analysis.

Similar results were observed when the analysis was restricted to those with a small aneurysm at baseline (growth= −0.10 mm/year [−0.23, 0.02] per copy of the minor allele; Supplementary Figure 4) or medium aneurysm at baseline (growth= −0.08 mm/year [−0.21, 0.05] per copy of the minor allele; Supplementary Figure 5). We did observe an association between the Asp358Ala variant and time to surgery threshold after adjusting for age and sex (hazards ratio (HR)= 0.85 [0.73, 0.98] per copy of the minor allele; Supplementary Figure 6a). The HR was in the same direction but became statistically non-significant in the subset of studies were we were able to additionally adjust for current smoking, diabetes status, body mass index and measurement method (HR=0.91 [0.77, 1.06]; Supplementary Figure 6b). Overall change in AAA growth remained the same when individuals with only a single measure of aneurysm size were included in the model (n=2,691, growth=-0.06 mm/year [− 0.18, 0.06]; Supplementary Figure 7).

### Inhibition of IL-6 signalling pathway in angiotensin II + anti-TGFβ mouse model

We next tested the effect of blocking the IL-6 pathway in two distinct, previously characterised mouse models of AAA (Methods). In the angiotensin II + anti-TGFβ model, mice infused with anti-IL-6R (blocking both classical and trans-signalling pathways) demonstrated a significant increase of plasma concentration of IL-6, as compared to isotype treated mice, and this difference was sustained over the course of the experiment (Figure 2A). We observed a reduction in plasma concentrations of serum amyloid A (SAA), a protein expressed in response to inflammation, after blocking IL-6R compared to the control mice (Figure 2B). Blocking IL-6R significantly reduced plasma concentration of IL-2 before and after the infusion and reduced concentration of IL-5 and chemokine ligand 1 (CXCL1) after the infusion (Figures 2A). After anti-IL-6R treatment, systolic blood pressure was significantly lower after the infusion compared to control treatment (Figure 2C). However, there was no significant difference in rate of aneurysm rupture between the anti-IL-6R treated and control groups (Figure 2D). Since there was no observed association with AAA rupture, we did not further assess the effect of blocking the IL-6R pathway on AAA growth.

**Figure 2:**
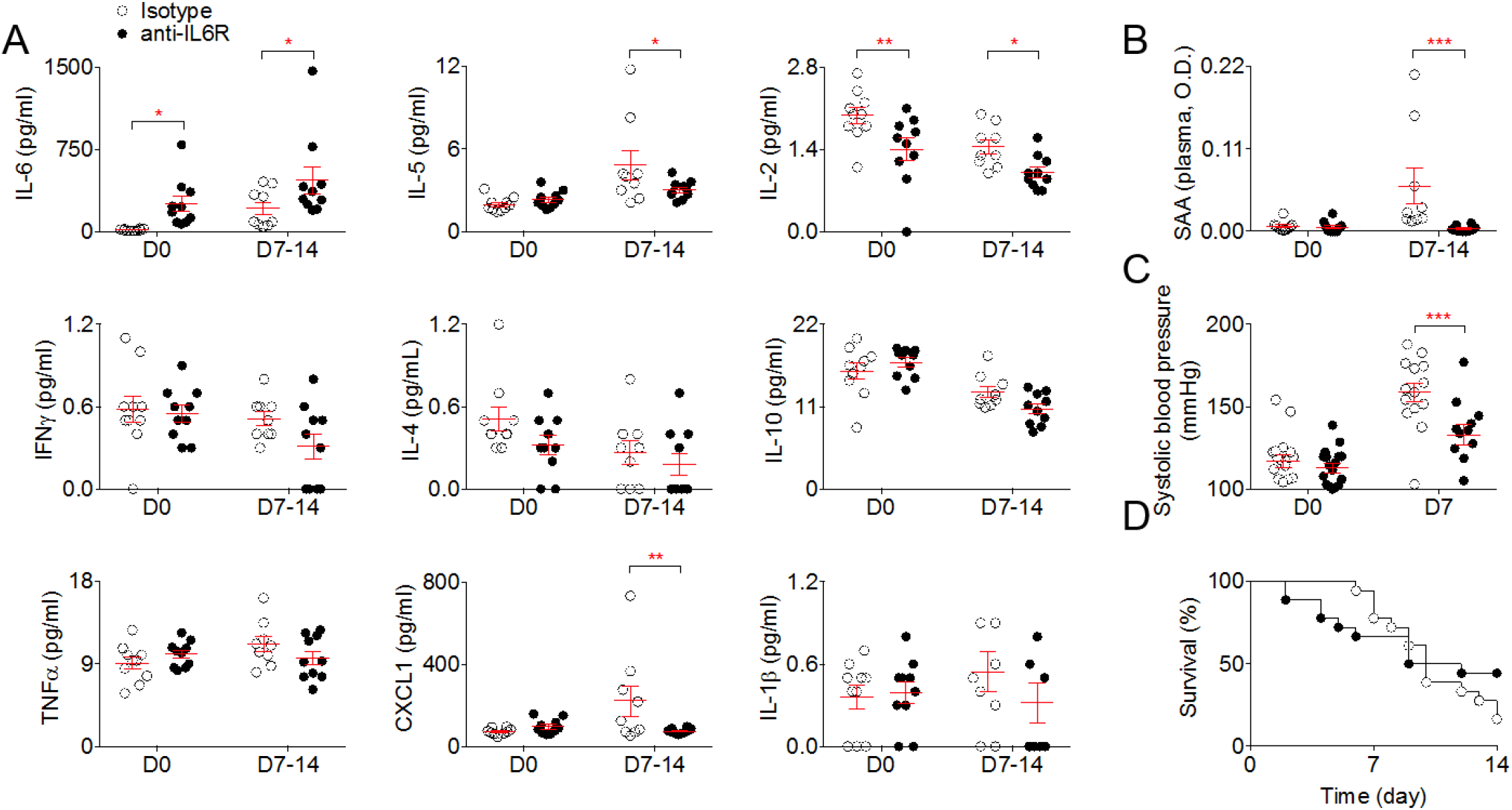
Anti-IL6-R prevents angiotensin II induced hypertension but does not protect against aortic rupture induced by angiotensin II and anti-TGFβ infusion. Mice were treated with anti-IL6-R or isotype control (n=22 mice/group) starting one week before angiotensin II and anti-TGFβ infusion. A - Plasma concentration of cytokines at day 0 (before angiotensin II and anti-TGFβ infusion) and day 7 to 14. *p<0.05 isotype vs anti-lL6-R; **p<0.01 isotype vs anti-lL-6R; 2-way ANOVA followed by uncorrected fisher’s test. B - Plasma concentration of serum amyloid A (SAA) at day 0 (before angiotensin II and anti-TGFβ infusion) and day 7 to 14. ***p<0.05 isotype vs anti-IL6-R; 2-way ANOVA followed by uncorrected fisher’s test. C - Systolic blood pressure measurement using tail cuff at day 0 and day 7 after angiotensin II and anti-TGFβ infusion. *‘*p<0.001 isotype vs anti-IL-6R; 2-way ANOVA followed by uncorrected fisher’s test. D - Survival curves of mice after angiotensin II and anti-TGFβ infusion. Results are coming from two independent experiments pooled together.

Selectively blocking the IL-6 trans-signalling pathway using sgp130Fc did not change the concentration of IL-6 (Figure 3A) or SAA (Figure 3B), but significantly induced IL-5 and reduced TNFα plasma concentration (Figure 3A). Although we observed no difference in systolic blood pressure between mice treated with sgp130Fc and the control mice (Figure 3C), there was a significant reduction in aneurysm rupture after sgp130Fc treatment compared to control treatment (Figure 3D).

**Figure 3:**
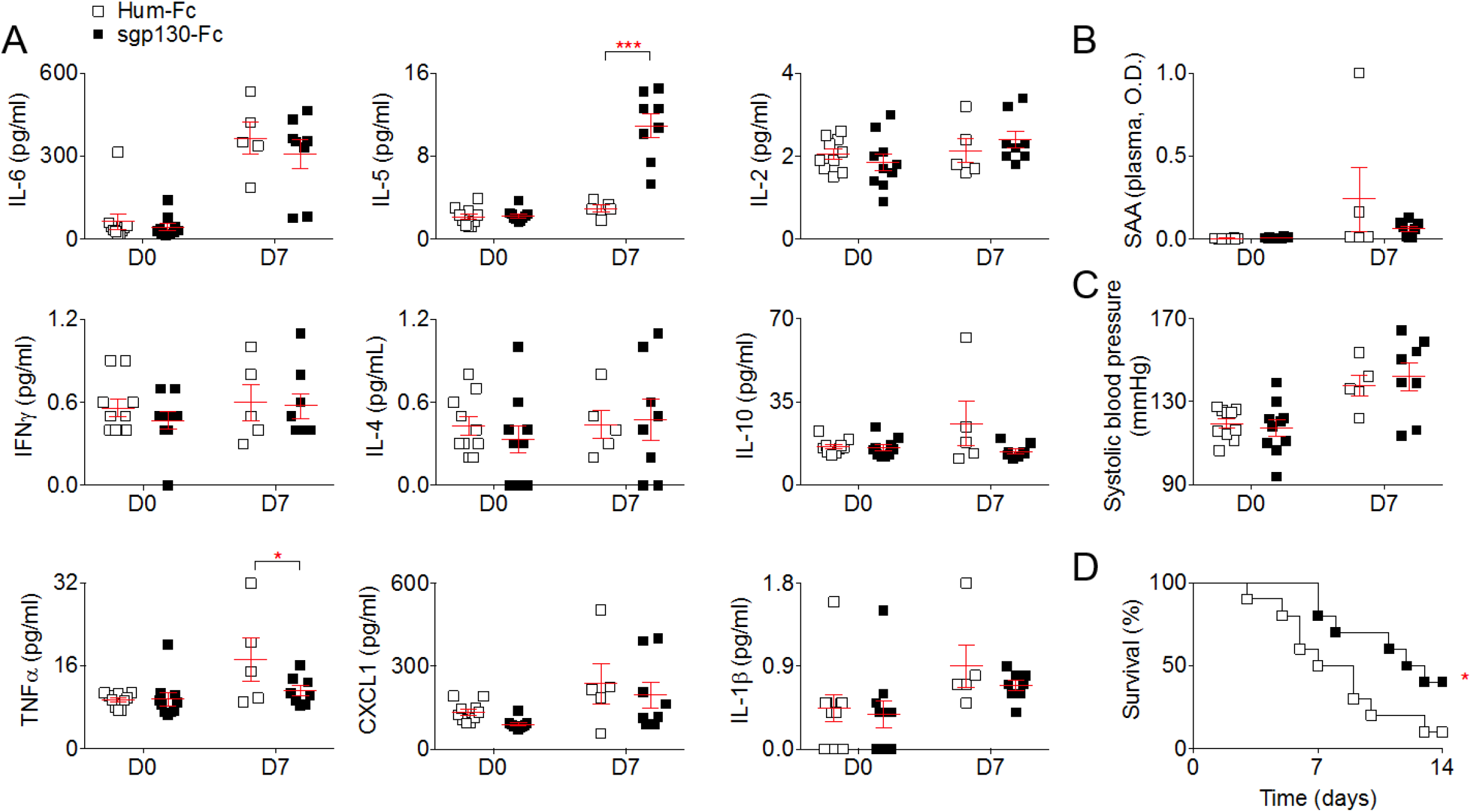
Selective blockage of the IL-6 trans-signalling pathway using sgp 130 in the angiotensin II+ anti-TGFβ model reduces aortic rupture. Mice were treated with sgp130-Fc or human lgG1 Fc (Hum-Fc) (n=10 mice/group) starting one week before angiotensin II and anti-TGFβ infusion. A - Plasma concentration of cytokines at day 0 (before angiotensin II and anti-TGFβ infusion) and day 7. *p<0.05 Hum-fc vs sgp 130; ***p<0.001 Hum-Fc vs sgp 130; 2-way ANOVA followed by uncorrected fisher’s test. B-plasma concentration of serum amyloid A (SAA) at day 0 (before angiotensin II and anti-TGFβ infusion) and day 7 C – Systolic blood pressure measurement using tail cut at day 0 and day 7 after angiotensin II and anti-TGFβ infusion. D – Survival curves of mice after angiotensin II and anti-TGFβ infusion. *p<0.05 Hum-Fc vs sgp 130; Gehan-Breslow-Wilcoxon test. Result are coming from one experiment.

### Inhibition of IL-6 signalling pathway in elastase + anti-TGFβ mouse model

Using the elastase + anti-TGFβ model, we found that blockage of the IL-6R pathway using anti-IL-6R resulted in significantly increased mortality (Figure 4A) induced by aortic rupture (Figure 4B) but there was no change in the diameter of the aneurysm at the end of the experiment (Figure 4C). There was also no change in the collagen content of the aortic wall (Figure 4D) or the recruitment of myeloperoxidase positive (MPO^+^) cells (Figure 4E), but treatment with anti-IL-6R significantly enhanced the recruitment of CD3^+^ T cells in the aortic wall (Figure 4F).

**Figure 4:**
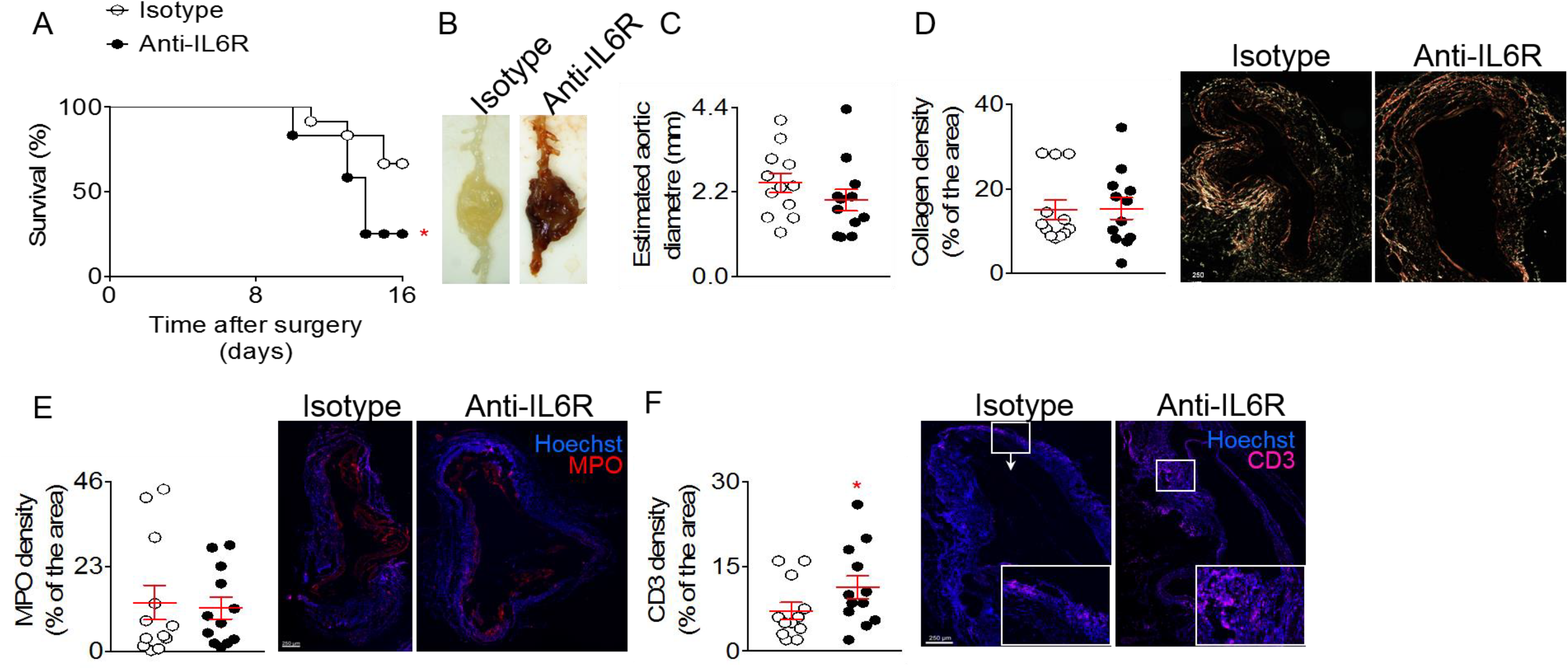
Blockage of the IL-6 pathway using anti-IL-6R in the elastase + anti-TNFβ model enhances T cell infiltration and rupture of the aorta. Mice were treated with anti-IL-6R or isotype control (n=12 mice/group) starting one week before the application of elastase and the infusion anti-TGFβ. A – Survival curves of mice after the application of elastase and the infusion anti-TGFβ. *p<0.05 isotype vs anti-IL6-R; Gehan-Breslow-Wilcoxon test. B – Representative macroscopic pictures of abdominal aortic aneurysms from mice treated with elastase and anti-TGFb and isotope or anti-IL6R, at day 16.Note that the aneurysm from the isotype treated mouse was not ruptured. C – Analysis of the aortic diameter (μm) based on the perimeter obtained from aortic cross sections. D – Quantification and representative images of collagen content of the aortic wall analysed using Sirius Red staining under polarized light. E, F – Quantification and representative images of MPO (D) and CD3 (E) immunofluorescent stainings on aortic cross section. *p<0.05 isotype vs anti-IL-6R; Mann-Whitney test. Results are coming from one experiment.

Blocking only the IL-6 trans-signalling pathway using sgp130Fc significantly increased survival (Figure 5A) by reducing aortic ruptures (Figure 5B), although at the end of the experiment there was no change in the aortic diameter between the treated and control mice (Figure 5C). Histological analysis of aortic samples revealed a significant increase in the collagen content of the arterial wall (Figure 5D) but no differences in Ly6G^+^ (Figure 5E) and CD3^+^ T cell recruitment (Figure 5F) after sgp130Fc infusion, as compared to the control mice. Table 2 summarises the results of the different mouse models with a comparison to the human genetic data.

**Figure 5:**
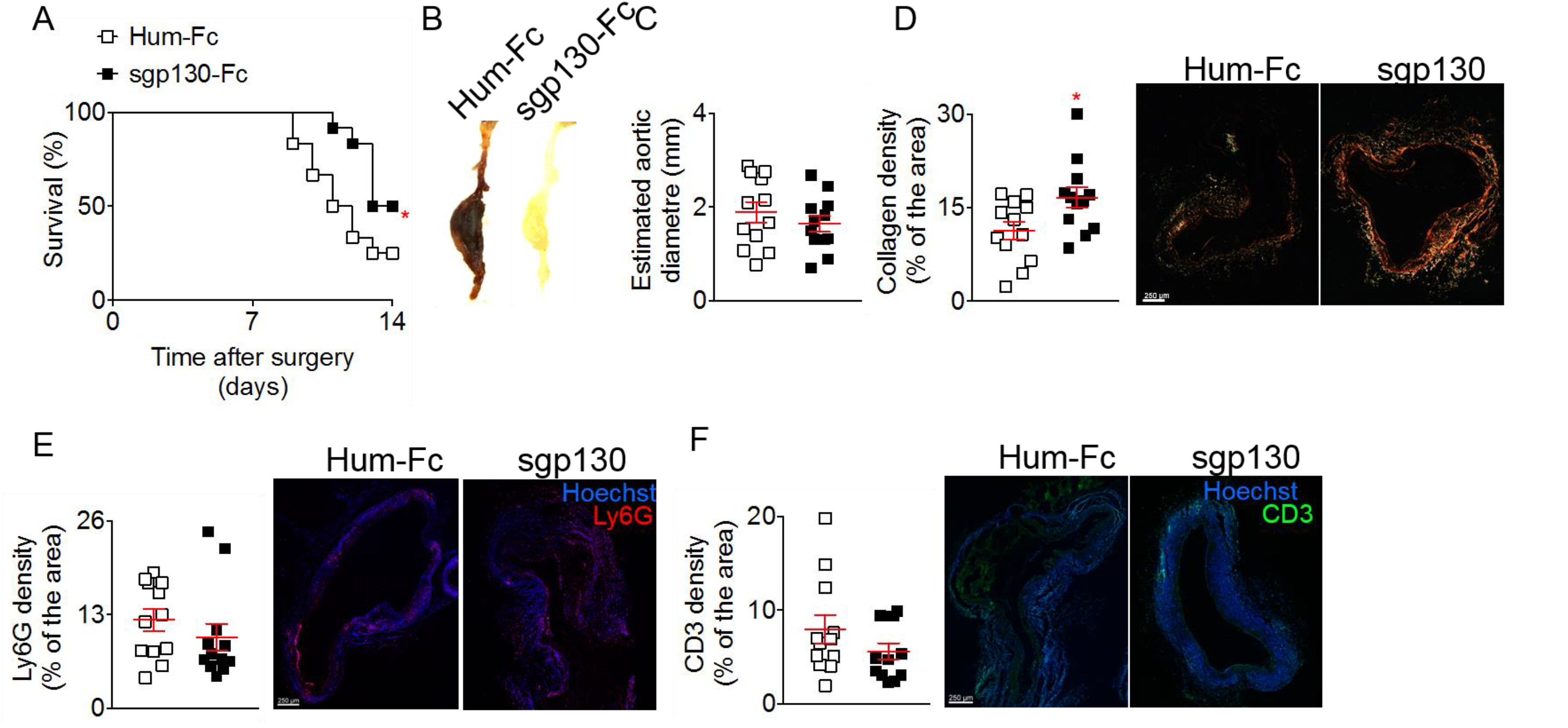
Selective Blockage of the IL-6 trans-signalling pathway using sgp 130 in the elastase + anti-TGFβ model increased collagen deposition and prevent aortic. Notes: Mice were treated with sgp 130 or Hum-Fc (n=12 mice/group) starting on the day of the application of elastase and the infusion anti-TGFβ. A - Survival curves of mice after the application of elastase and the infusion anti-TGFβ. *p<0.05 Hum-Fc vs sgp130; Gehan-Breslow-Wilcoxon test. B - Representative macroscopic pictures of abdominal aortic aneurysms from mice treated with elastase and anti-TGFb and isotope or anti-IL6R, at day 16.Note that the aneurysm from the isotype treated mouse was not ruptured. C - Analysis of the aortic diameter (μm) based on the perimeter obtained from aortic cross sections. D - Quantification and representative images of collagen content of the aortic wall analysed using Sirius Red staining under polarized light. *p<0.05 Hu-Fc vs sgp 130; Mann-Whitney test. E, F - Quantification and representative images of Ly6G (D) and CD3 (E) immunofluorescent staining on aortic cross section. Result are coming from one experiment.

**Table 2.** Comparison of results for the association between IL-6R and abdominal aortic aneurysm from human genetic analysis and mouse experimental models.

| **Model** | **Inhibition of IL-6 signalling** | **Effect on AAA growth/ survival** | **p-value of effect** | **Other cardiovascular markers (p<0.05)** |
| --- | --- | --- | --- | --- |
| **Humans stratified for *IL6R*-Asp358Ala** | Minor allele associated with dampening of classical signalling. Effects on trans-signalling not clear. | Non-significant change in AAA growth (0.06mm/year)* | 0.136 | Increased IL-6, sIL-6R, CXCL10, CXCL11, IL1α and CD6;<br>Reduced MMP3, OPG, IGFBP1, TIMP4 |
| **Angiotensin II + anti-TGFβ mouse model** | Both classical and trans-signalling | No difference in survival | 1.000 | Increased IL-6;<br>Reduced SAA, IL-2, IL-5, CXCL1 and SBP |
| | Only trans-signalling | Improved survival | 0.040 | Increased IL-5;<br>Reduced TNFα |
| **Elastase + anti-TGFβ mouse model** | Both classical and trans-signalling | Decreased survival | 0.029 | Increased recruitment of CD3<sup>+</sup> T cells |
|  | Only trans-signalling | Improved survival | 0.024 | Increased collagen content of the arterial wall |

\*Effect indicated per copy of the IL6-R minor allele.

### Evaluation of the effect of IL6R-rs2228145 on a range of cardiovascular markers

As would be expected, the minor allele of *IL6R*-rs2228145 was associated with increased plasma concentrations of IL-6 and sIL-6R. The variant was also associated with increased monocyte count, after correcting for multiple comparisons (p<1.389×10^-3^). At a nominal significance level (p<0.05), rs2228145-C was associated with reduced lymphocyte count, increased levels of the cytokines CXCL10, CXCL11 and IL1α, as well as CD6 (an important regulator of T cells), and reduced levels of MMP3 (a matrix metalloproteinase), TIMP4 (a metalloproteinase inhibitor), OPG (osteoprotegerin) and IGFBP1 (Supplementary Figure 8). In our analysis, the effects on plasma IL-2, IL-5, and CXCL1 levels, as well as on blood pressure, were not statistically significant.

## Discussion

In a combined analysis of the available worldwide clinical genetic data on AAA growth, we observed no statistically significant decrease in annual AAA growth rates for carriers of the minor allele of the Asp358Ala variant (rs2228145) in the *IL6R* gene. While we did observe a 15% decrease in the rate of reaching the surgery threshold of ≥55mm (HR=0.85 [0.73, 0.98] per copy of the minor allele), people with copies of the *IL6R*-Asp358Ala variant also had, on average, smaller baseline aneurysm diameters. Although we tried to account for this by allowing baseline hazards to vary depending on initial aneurysm size, some residual confounding is possible and could explain the observed results. In experimental data from mouse models, we found that selective blockade of the IL-6 trans-signalling pathway was associated with decreased aortic rupture and death. In exploratory analyses of cardiovascular and inflammatory biomarkers in healthy participants, we found that rs2228145-C was inversely associated with plasma levels of osteoprotegerin, matrix metalloproteinase-3 and metalloproteinase inhibitor-4 (p<0.05). Osteoprotegerin has previously been shown to promote matrix metalloprotease release from monocytes and vascular smooth muscle cells,^34, 35^ and aberrant aortic extracellular matrix remodelling has been suggested to play a key role in the pathogenesis of AAA.^36^ However, we note that further studies are needed to validate our biomarker data. Taken together, these human genetic, biomarker, and experimental murine findings are compatible with the concept that IL-6 trans-signalling is relevant to AAA growth, encouraging larger-scale evaluation of this hypothesis.

There are two potential pathways through which IL-6 signalling may affect disease development and progression: classic signalling and trans-signalling (Figure 6). While the minor allele [C] in rs2228145 dampens classic signalling, the effect of the allele on trans-signalling has not conclusively been established. As trans-signalling effects are likely to depend on local availability of IL-6, sIL-6R and sgp130, they are context-specific and may differ across tissues. The sgp130-sIL-6R complex acts as a “buffer” for IL-6, regulating the concentration of available active IL-6, for example, during inflammation. The capacity of the buffer is determined by the levels of sIL-6R, as these are lower than the levels of sgp130.^14^ If increased availability of sIL-6R results in a dampening of the IL-6 trans-signalling pathway, this may explain potential protective effects in AAA and is consistent with previously observed protective effects in mouse models of sepsis^37^ and pancreatic and lung failure.^38^ As we found a consistent pattern of results when the trans-signalling pathway was selectively blocked in the mouse models, it suggests that this pathway could have a detrimental effect on AAA growth (Figure 6). For example, the minor allele of the rs2228145 variant may result in a local reduction of IL-6 trans-signalling in the abdominal vasculature, reducing AAA risk^12^ and, perhaps, AAA growth rates. Our observation of only a small, but statistically insignificant, decrease in annual AAA growth rates does not preclude meaningful clinical effects, since the growth rate reduction estimated from natural genetic variation does not necessarily relate to the magnitude of the benefit that might result from pharmacological treatment directed at the IL-6 trans-signalling pathway.^39^ Selective blocking of the IL-6 trans-signalling pathway using sgp130Fc is being investigated in phase II clinical trials in patients with inflammatory bowel disease.^14^

**Figure 6:**
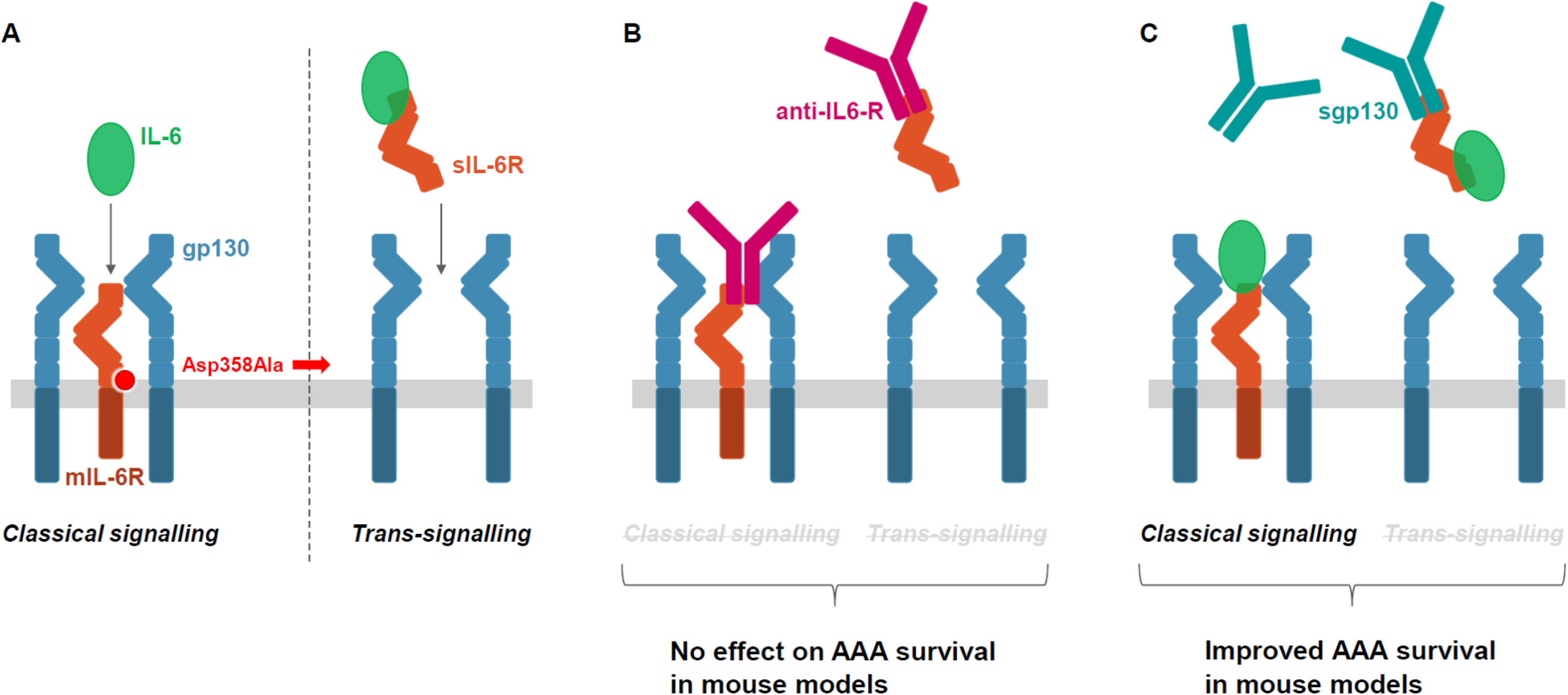
Overview of the IL-6 classical and trans-signalling pathways and their potential role in AAA growth. Notes: In “classical” IL-6 signalling (left), the binding of the cytokine IL-6 to the membrane-bound IL-6 receptor (mIL-6R) leads to the dimerization of its co-receptor gp 130, and subsequently. triggers downstream signalling in a restricted subset of cells. In IL-6 trans-signalling (right), IL-6 forms a complex with soluble IL-6 receptor (sIL-6R) that can stimulate cells expressing gp 130 even in the absence of mIL-6R. The minor allele of a functional variant in the *IL6R* gene, Asp358Ala (rs2228145A>C) results in more efficient proteolytic cleavage of mIL-6R, thereby reducing levels of mIL-6R and classical IL-6 signalling but potentially increasing trans-signalling. B – In mouse models AAA, the blockage of both the classical and trans-signalling pathways with anti-IL6-R (i.e. MR16-1, the animal-equivalent of tocilizumab) did not have a conclusive effects on the time to aneurysm rupture. C – Specific blockage of the trans-signalling pathway with with sgp 130 resulted in improved survival rates in mouse of AAA.

Nevertheless, apparent inconsistencies in our findings require further elucidation. For example, blockade of both the classic and trans-signalling IL-6 pathways using the animal-equivalent (MR16-1) of tocilizumab had no effect on AAA rupture in the angiotensin II + anti-TGFβ model, but it was associated with decreased survival in the elastase + anti-TGFβ model. These different outcomes could be explained by differences in the development of AAA in the different mouse models. The primary process of “aneurysm” formation in the angiotensin model is a medial dissection, which may be accentuated by elevations in blood pressure (even though high blood pressure is not the primary cause of medial dissection). Hence, the potential protection afforded by MR16-1 antibody in this model can at least in part be attributed to the significant reduction of blood pressure. In contrast, the elastase model does not involve medial dissection or elevations in blood pressure, but induces progressive remodelling, dilatation, and eventually transmural rupture of the artery wall, better mimicking AAA progression in humans.^29, 30^

If selective blockade of the IL-6 trans-signalling pathway results in decreased aortic rupture, as suggested by our murine data, one might expect that blocking both the classic and trans-signalling pathways would also result in decreased aortic rupture. However, we did not observe such a finding, perhaps due to competing downstream actions of the classic and trans-signalling pathways. Such an explanation is consistent with our finding that IL-5 levels were increased and TNFα levels decreased when trans-signalling was selectively blocked, whereas blockade of both classic and trans-signalling pathways led to reductions in IL-5 levels and no changes in TNFα. Our findings suggest that selective blockade of the IL-6 trans-signalling pathway, compared to blockade of both IL-6 signalling pathways, results in different downstream cytokine profiles and potentially different effects on AAA progression. It is also possible that the blocking of both the classical signaling cascade (considered to have protective and regenerative cellular effects) and trans-signaling cascade (considered to have pro-inflammatory effects) cancelled each other out, leading to no detectable effect on AAA rupture.

We undertook a range of sensitivity analyses to test assumptions underlying our longitudinal human genetic studies. We studied complementary murine models of AAA, including the elastase + anti-TGFβ mouse model that has been shown to more closely mimic the AAA growth and rupture patterns seen in humans. To generate new mechanistic hypotheses, we conducted exploratory studies of the *IL6R*-Asp358Ala variant in relation to cardiovascular and inflammatory plasma biomarkers recorded in healthy participants. The experimental mouse studies were conducted under severe conditions in which TGFβ was blocked. Although any effect of IL-6R signalling might have been easier to observe in a less severe model, if the intervention is protective in this more severe model it provides assurance that the intervention will also be protective in a less severe model.

Our study had potential limitations. It was powered to detect reductions in aneurysm growth of ~0.21mm per year or larger, much greater than the observed non-significant decrease of 0.06mm per year. Future studies powered to see an effect the same size as that observed in the current study would need to recruit an additional ~21,500 participants (total participants needed=24,444, Appendix 3). This is unlikely to be achievable in the near future; alternative study methods using a composite phenotype for disease progression may be needed. Index event and survival bias, in which participants are selected into the study based on both having and surviving an event, may have biased the results towards the null.^40^ However, this bias is likely to be small (<10%). Further, ultrasound, the primary method used to assess AAA diameter in the included studies, has a margin of error of 2-3mm,^41^ greater than the annual rate of aneurysm growth, making changes in growth difficult to detect. This might be why we observed an association between the *IL6R*-Asp358Ala variant and time to surgery threshold of ≥55mm but not when looking at continuous change in AAA size. Although we examined rupture rather than aortic diameter as the outcome in the mouse experimental models, our published data indicate that aortas that rupture have larger diameters or faster diameter progression than the ones that do not rupture.^30^

It is also uncertain how well the results of our animal models translate to clinical disease. For example, an important difference is that IL-6 blockage is initiated before or at the time of disease development in the mice models of AAA, thereby not truly mimicking the treatment effects expected in humans, in which drugs to block IL-6 pathway would be started after disease onset. Blockade of the IL-6 signalling pathways in the angiotensin II + anti-TGFβ mouse model resulted in reproducible reductions in systolic blood pressure. Although tocilizumab has been anecdotally reported to improve pulmonary hypertension in Castleman’s disease,^42^, 43 the rs2228145 variant was not associated with changes in systemic blood pressure in healthy participants in a genome-wide association study. A large-scale randomized trial found no difference in the number of hypertension events reported in those using tocilizumab compared to placebo.^27, 44^ Thus, the acute responses to pharmacological doses of angiotensin II in the mouse model may not faithfully reproduce the human setting of AAA.

Our study may have clinical implications. Tocilizumab is currently indicated in a few disease settings, including rheumatoid arthritis and giant cell arteritis, both of which are associated with an increased risk of aortic aneurysm.^45, 46^ The development of coronary artery aneurysms has also been reported in a non-placebo-controlled pilot study of tocilizumab in children with Kawasaki’s Disease.^47^ Our finding that blocking both the classic and trans-signalling IL-6 pathways using the animal-equivalent of tocilizumab was associated with decreased survival in the elastase + anti-TGFβ model provides some support that aortic aneurysm progression should be monitored in these patients.

In conclusion, our proof-of-principle data are potentially compatible with the concept that IL-6 trans-signalling is relevant to AAA growth, encouraging larger-scale evaluation of this hypothesis.

## Acknowledgments

We thank Chugai Pharmaceutical Co., Ltd. for providing the MR16-1. We thank Tao Jiang and Praveen Surendran (Cardiovascular Epidemiology Unit, University of Cambridge) for providing access to genomic datasets.

## Sources of Funding

The Cardiovascular Epidemiology Unit is supported by the UK Medical Research Council (MR/L003120/1), British Heart Foundation (RG/13/13/30194), and NIHR Cambridge Biomedical Research Centre. SRJ was supported by grants of the Deutsche Forschungsgemeinschaft (CRC877, project A1) and the German Cluster of Excellence 306 ‘Inflammation at Interfaces’.

## Disclosures

Since October 2015, Daniel F. Freitag has been a full time employee of Bayer AG, Germany.

